# Detection of antibodies against a conserved capsid epitope as the basis of a novel universal serological test for foot-and-mouth disease

**DOI:** 10.1101/797332

**Authors:** A Asfor, N Howe, S Grazioli, S Berryman, K Parekh, G Wilsden, A Ludi, DP King, S Parida, E Brocchi, TJ Tuthill

## Abstract

Diagnostic tests for foot-and-mouth disease (FMD) include the detection of antibodies against either the viral non-structural proteins or the capsid. The detection of antibodies against the structural proteins (SP) of the capsid can be used to monitor seroconversion in both infected and vaccinated animals. However, SP tests need to be tailored to the individual FMD virus serotype and their sensitivity performances may be affected by antigenic variability within each serotype and mismatching between tests reagents. As a consequence, FMD Reference Laboratories need to maintain contingency to employ multiple type-specific assays for large-scale serological surveillance and post-vaccination monitoring in the event of FMD outbreaks. In this study, a highly conserved region in the N terminus of FMDV capsid protein VP2 (VP2N) was characterised using a panel of intertypic-reactive monoclonal antibodies. This revealed a universal epitope in VP2N which could be used as a peptide antigen to detect FMDV-specific antibodies against all types of the virus. A VP2-peptide ELISA (VP2-ELISA) was optimised using experimental and reference antisera from immunized, convalescent and negative animals (n=172). The VP2-ELISA is universal, simple and provided sensitive (98.6 %) and specific (93%) detection of antibodies to all FMDV strains used in this study. We anticipate that this SP test could have utility for sero-surveillance during virus incursions in FMD-free countries and as an additional screening tool to assess FMD virus circulation in endemic countries.

## Introduction

Foot-and-mouth disease (FMD) is an economically devastating viral disease of cloven-hoofed animals with a global distribution. It limits access to markets for developing countries and outbreaks in otherwise FMD-free countries are expensive to control (as in the UK in 2001, Japan in 2010 and the Republic of Korea in 2010 and 2011) [1, 2]. FMD virus (FMDV) is a single-stranded, positive-sense, RNA virus belonging to the genus *Aphthovirus* in the family *Picornaviridae*. The virus exists as seven serotypes (O, A, C, Asia 1, South African Territories (SAT)1, SAT2 and SAT3) as well as numerous and constantly evolving strains showing a spectrum of antigenic diversity.

The non-enveloped picornavirus capsid has icosahedral symmetry, a diameter of approximately 30 nm and is composed of 60 copies of each of the capsid proteins VP1, VP2, VP3 and VP4. VP1, VP2 and VP3 are the major components of the capsid, while VP4 is a small (approximately (12 kDa) internal protein which lies on the inside surface of the capsid around the five-fold axes of symmetry, where it is thought to stabilise interactions between pentameric capsid subunits [3, 4]. During the replication cycle of FMDV, eight different viral non-structural proteins (NSPs; and additional precursors) are generated which are potential serological targets for diagnostic assays [5]. The presence of antibodies against NSPs can be used to differentiate infected and vaccinated animals (DIVA) because such antibodies are only produced by infection and are not elicited after administration with purified vaccines. In addition, the inter-serotypic conservation of the NSPs means this type of test is compatible with all serotypes of FMDV. Hence, NSP tests can be used as generic screening tools to support national programs to attain the OIE status of FMD-freedom with or without vaccination [6, 7, 8]. However, the specificity of these tests is less than 100% [9] and testing algorithms that are designed to confirm absence of FMDV circulation in large populations usually adopt screening and confirmatory serological assays with covariant rates of false positivity [7, 8, 9]. In this context, ELISAs that measure FMDV-specific antibodies directed at capsid structural proteins (SP) are widely used to augment NSP tests for sero-surveillance activities [10, 11, 12, 13]. One of the international standard tests for FMDV antibody detection is the virus neutralisation test (VNT) [14]. However, the VNT is laborious, rendering large scale serological testing difficult. In addition, the procedure requires live virus, thus confining the test to high containment laboratories in non-endemic countries. SP ELISAs with high diagnostic sensitivity are also available for certification of animals as free from FMD prior to import and export, for serological confirmation of FMDV infection, for post vaccination monitoring and for the demonstration of vaccine efficacy [14]. However, SP assays need to be tailored to individual serotypes and as a consequence FMD Reference Laboratories need to maintain parallel assay systems to accommodate the possibility of FMD outbreak due to different virus serotypes.

A number of monoclonal antibodies (mAbs) have previously been reported with cross-reactivity against multiple FMDV serotypes [15, 16, 17]. The recognition sites for some of these mAbs have been mapped to a highly conserved region at the N-terminus of VP2 [15, 16, 17]. In this study, a highly conserved region in the N terminus of FMDV capsid protein VP2 (VP2N) was characterised using a panel of cross-reactive mAbs. This revealed a universal epitope in VP2N which has been investigated as a peptide antigen to detect FMDV-specific antibodies in serum samples from animals infected or vaccinated with any of the FMDV serotypes.

## Material and Methods

### Cells lines and Viruses

The IBRS-2 (pig kidney) cell line and the BHK-21 (baby hamster kidney 21) cell line, used for FMD viruses propagation and immunoassays, were maintained either in Dulbecco’s modified Eagle’s medium or in Minimum Essential Medium, (DMEM; Thermo-Fisher Scientific, UK) supplemented with 10% heat-inactivated foetal bovine serum (FBS; Thermo-Fisher Scientific, UK) and 100 U of penicillin-streptomycin (Sigma) per ml. FMDV strains used are indicated in each relevant paragraph.

### Peptides

Peptides representing the N-terminal 15 (VP2N15), 30 (VP2N30) or 45 (VP2N45) amino acids of FMDV VP2 were synthesised (Peptide Protein Research, UK) without modifications except for the addition of 6 lysines at the C-terminus of the peptides to increase the solubility. VP2N45 was used for the development of the peptide ELISA. A control peptide equivalent to a capsid sequence from the related picornavirus human rhinovirus was used [18]. Eight peptides (15mer each) overlapping by ten amino acids, covering the first 45 amino acids from the N-terminus of the FMDV capsid sequence, were used for the fine mapping of the epitope (Fig.1a).

**Fig.1.**
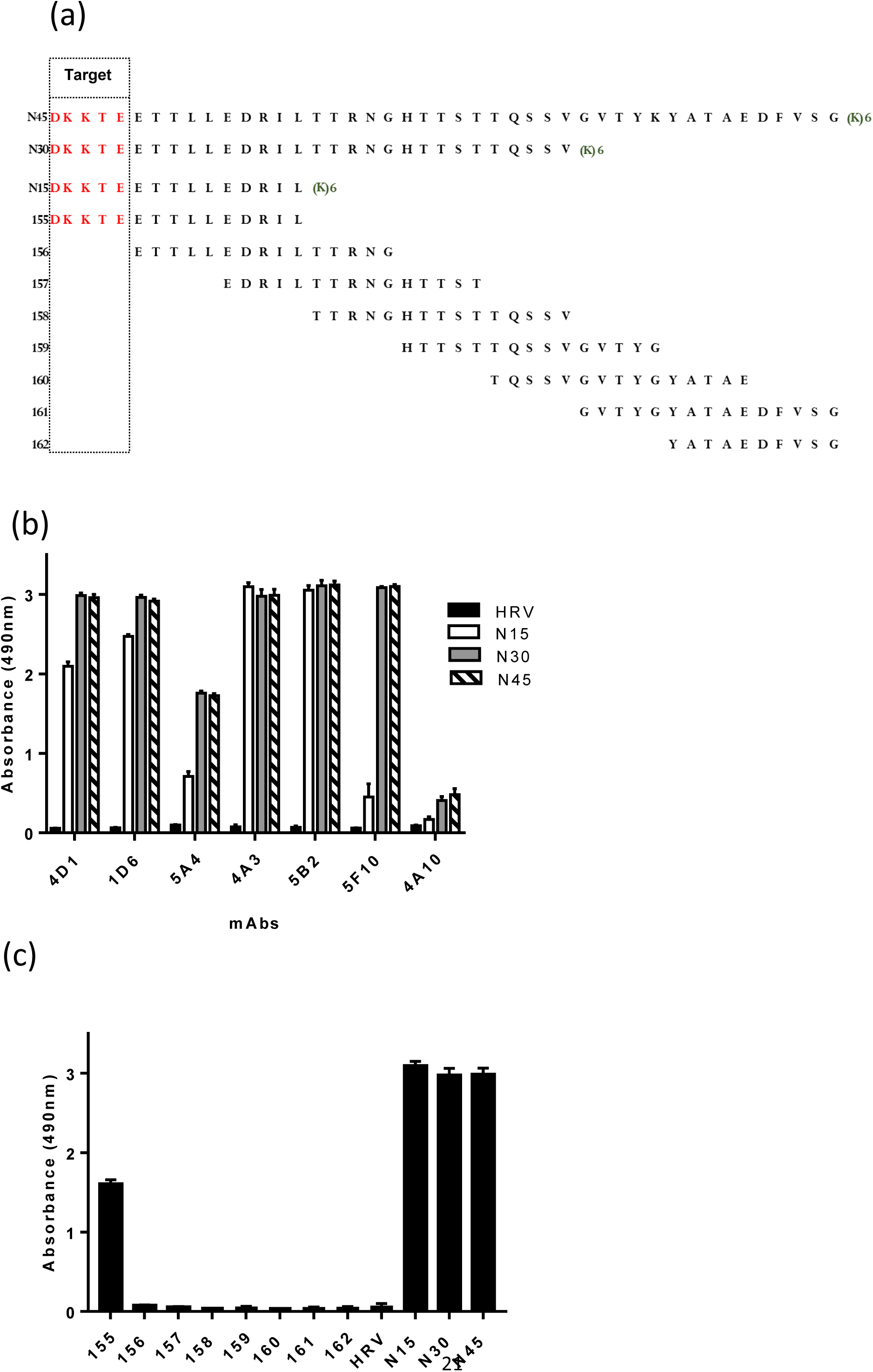
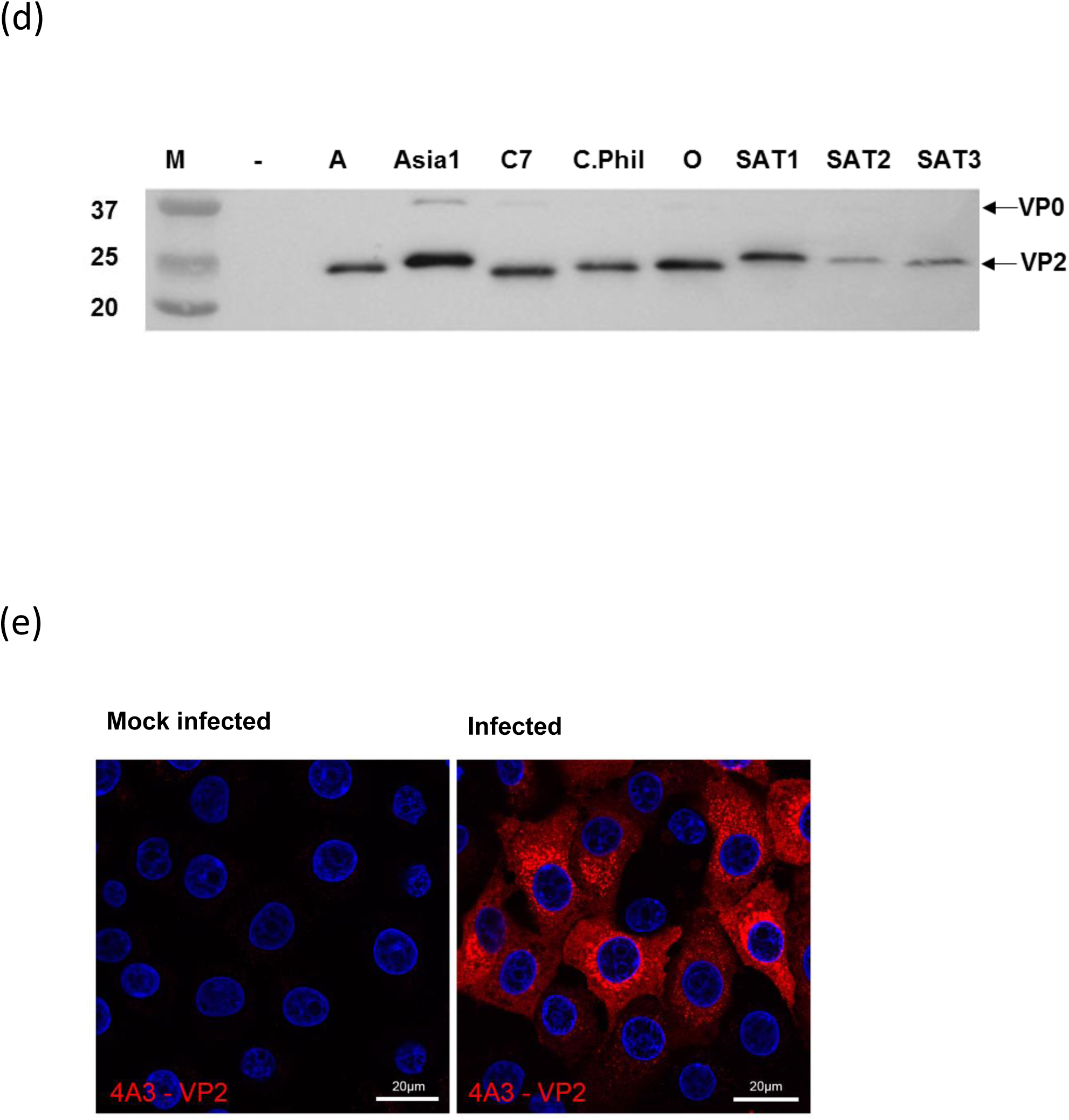
FMDV heterotypic-reactive mAbs recognise the N terminus of VP2. **(a)** Overlapping peptides representing the VP2 N-terminal 45 amino acids. The (K)6 denotes to addition of 6 lysine residues at the C-terminus of the peptide to increase peptide solubility. **(b)** Peptide ELISA showing cross reactive mAbs recognise peptides equivalent to the N-terminal 15 (N15), 30 (N30) and 45 (N45) amino acids of FMDV VP2. The N-terminal 45 amino acids of human rhinovirus VP4 (HRV-VP4) was used as negative control; peptides concentration was 2 µg/ml. **(c)** mAb 4A3 epitope mapping (using peptides shown in panel a) identifies the cross-reactive epitope at the N-terminus of VP2. **(d)** Reactivity of mAb 4A3 with capsid protein VP2 of all 7 serotypes in western blot. 4A3 mAb produced a clear intense band for VP2 and a weaker reaction for VP0 **(e)** Immunofluorescence microscopy using mAb 4A3 to detect FMDV serotype O infected IBRS-2 cells.

### Serum samples

Sera from infected cattle with FMDV O/UKG 34/2001 [19] was used to optimise the ELISA. Reference sera from experimentally vaccinated or infected animals were supplied by FAO World Reference Laboratory for FMD (WRLFMD, The Pirbright Institute). The parameters of selecting serum samples were as follows: Negative (n=100): samples that been collected from negative coherent country (during the UK 2007 outbreak). These samples are from non-vaccinated animals and proved to be negative using NSP-ELISA. Positive (n=72): samples that are known to be infected or vaccinated with FMDV. Selection of the positive samples was based up on more than 7days post vaccination or infection to ensure a positive response. See supplementary table (1) for more details.

### Production of mAbs

The following FMD viruses were used as immunogens to produce mAbs in mice and for the following selection of heterotypic cross-reactive mAbs: serotype A Malaysia 16/97, C1 Brescia 1964, Asia 1 Nepal 29/97, A24 Cruzeiro and O UK 31/2001.

For each immunogen, BALB/c mice were primed subcutaneously with 20μg of purified FMD virus in Freund’s complete adjuvant and boosted intraperitoneally with the same antigen in phosphate buffered saline (PBS) once or twice at one-month intervals. Three days after the last boost, mice were humanely sacrificed and hybridomas were generated by fusion of splenocytes with NS0 myeloma cells following standardized procedures [20]. Briefly, at least 10^8^ spleen cells were recovered from each mouse and fused with NS0 myeloma cells at a 10:1 ratio using PEG 4000. Fused cells diluted in Dulbecco’s modified Eagle medium, supplemented with hypoxanthine/aminopterin/thymidine and 20% fetal calf serum, were distributed over five microplates (200μl per well). Growing colonies were observed in all wells; in order to select hybridomas secreting monoclonal antibodies specific for the immunogen, the supernatants were screened by trapping ELISAs against the homologous virus strains. Selection of the inter-types cross reactive mAbs was based on results of the trapping ELISA against the homologous and heterologous virus serotypes, as previously described [21]. The selected hybridoma cells were cloned by limiting dilution in order to obtain antibodies from one single cell. The supernatant from exhausted cultures was then used as source of mAb.

### Immunofluorescence confocal microscopy

IBRS-2 cells on 13-mm glass coverslips (VWR) were infected with FMDV type O1 Kaufbeuren (MOI = 2) for 3.75 hours and then washed with PBS and fixed with 4% paraformaldehyde for 40 min at room temperature (RT). The cells were then permeabilized for 20 min with 0.1% Triton X-100 prepared in blocking buffer (Tris-buffered saline supplemented with 1 mM CaCl_2_, 0.5 mM MgCl_2_, 10% normal goat serum, and 1% fish skin gelatin). The cells were then incubated with primary antibody (mouse mAb 4A3) diluted 1/1000 in blocking buffer for 1h at RT. Subsequently, the cells were washed and incubated with Alexa-Fluor-conjugated secondary antibody (goat anti-mouse IgG Alexa-568; Thermo Fisher Scientific, UK) in blocking buffer for 45 min at RT. After washing, the cells were mounted using Vectashield mounting medium with DAPI (4,6-diamidino-2-phenylindole) (Vector Labs) and the coverslips sealed with nail varnish. All data were collected sequentially using a Leica SP8 confocal laser scanning microscope.

### SDS-PAGE and western blot

Initial tests to verify the reactivity in western blot of each mAb with the homologous partially purified strain were performed as previously described [21]. Later on, the cross-reactivity of one representative mAb (4A3) with all FMDV serotypes was confirmed as follows.

Virus lysates from IBRS-2 cells infected cells with different FMDV serotypes were denatured and reduced by heating at 95°C for 5min in Red Loading Buffer and DTT (NEB). The samples were resolved through 12% Tris-glycine gels and transferred to nitrocellulose membrane (0.45μM, GE Healthcare) using a Mini-Protean tetra cell (BioRad). Membranes were placed in blocking buffer (20mM Tris, 150mM NaCl pH7.6 with 0.1% v/v tween-20 (TBS-T) with 1% bovine serum albumin (BSA) w/v (Melford)) for 1h at RT followed by incubation with hybridomas supernatants (mAbs) and anti-mouse HRP-conjugated secondary antibody (Dako) (1/5000 in blocking buffer) in sequence for 1h at RT. Each incubation was separated by cycles of three washings with TBS-T. West Pico chemiluminescent substrate (Thermo Fisher Scientific, UK) was added to the membrane and exposures of the membrane were collected and visualised using a G: Box Chemi XX6 (Syngene).

### Serological standard tests: virus neutralisation test (VNT), liquid-phase blocking ELISA (LPBE), solid-phase competition ELISA (SPCE) and commercial kits (PrioCHECKTM FMDV Type O, Type A and Type Asia 1 Antibody ELISA kits)

VNT was carried out in microplates against 100 TCID_50_ of the homologous or heterologous viruses and results were reported as the final dilution required to neutralize 50% of the inoculated cultures [14]. The LPBE and the SPCE were carried out as described by Hamblin *et al*., (1986) [12] and by Paiba *et al*., (2014) [13] respectively. The cut offs used in the VNT (log titre 1.65), LPBE (log titre 1.95) and SPCE (40% of inhibition) were according to the standard operating procedures for the WRLFMD (The Pirbright Institute, UK). PrioCHECK ELISAs for FMDV type O, A and Asia 1 antibody were carried out according to the kits instructions, with 50% of inhibition as cut-offs.

The frequency distribution of values generated by various serological assays for the negative and the positive (vaccinated and infected animals) serum samples were plotted using GraphPad Prism (V7). Statistical analysis was performed using GraphPad Prism V7 for Windows (GraphPad Software, La Jolla California USA, www.graphpad.com).

### Indirect ELISAs and the development of the VP2 ELISA

Plastic 96-well plates (Maxisorp –Nunc) were coated with 100μl per well of the peptides in 0.05M standard carbonate/bicarbonate coating buffer (pH 9.6) at 4°C overnight. Different peptides concentrations, ranging from 125ng/ml up to 4µg/ml, were initially evaluated for test optimization. Wells were washed three times with phosphate buffered saline (PBS) containing 0.1% Tween 20 (PBS-T) between all incubations. Wells were blocked with 200μl blocking buffer (1% w/v BSA in PBS-T) at 37°C for 1h, and incubated either with 100μl of mAb (hybridoma supernatants, 1/5) or bovine sera (diluted 1:50 to 1 in 400 in blocking buffer) at 37°C for 1h. Antibody binding was detected by incubation at 37°C for 1h with 100μl of species specific HRP conjugated secondary antibodies (Dako), diluted in blocking buffer 1:1,000 in case of anti-mouse Ig conjugate or 1:15,000 for the anti-bovine-Ig conjugate. The chromogen development was mediated by the addition of 50μl of HRP substrate (OPD: Sigma FAST, Sigma, UK). The reaction was stopped after 20min by addition of 50μl of 1.25M sulphuric acid and the optical density (OD) was measured at 490nm.

## Results

### Characterisation of an FMDV-VP2 conserved epitope by cross reactive mAbs

Among the multiplicity of mAbs generated from mice independently immunized with four different FMDV serotypes (A Malaysia 16/97, C1 Brescia 1964, Asia 1 Nepal 29/97, A24 Cruzeiro, or O UK 31/2001), seven mAbs were selected because of their cross-reactivity with the seven FMDV serotypes. All mAbs were characterised as non-neutralising. Five of these mAbs strongly recognised the capsid protein VP2 by western blot and showed a weaker reaction with VP0, while two mAbs reacted with P1 (Table 1).

**Table 1.** FMDV mAbs showing cross-serotype reactivity and viral protein (VP) specificity.

| mAb ID | Parent virus | Trapping ELISA |  |  |  |  |  |  | Western blot |
| --- | --- | --- | --- | --- | --- | --- | --- | --- | --- |
|  |  | O | A | C | ASIA1 | SAT1 | SAT2 | SAT3 | VP TARGET |
| 5B2 | A Malaysia 16/97 | + | + | + | + | + | + | + | VP2 (+VP0) |
| 4A3 | C1 | + | + | + | + | + | + | + | VP2 (±VP0) |
| 5F10 | Asia 1 Nepal 29/97 | + | + | + | + | +/- | + | + | VP2 (+VP0) |
| 4A10 | A24 Cruz | + | + | + | + | + | + | + | P1 |
| 5A4 | A24 Cruz | + | + | + | + | + | + | + | P1 |
| 4D1 | O UK 31/01 | + | + | + | + | + | + | + | VP2 (+VP0) |
| 1D6 | O UK 31/01 | + | + | + | + | + | + | + | VP2 (+VP0) |

Previous studies have identified the conserved N-terminus of VP2 as a site for recognition by cross-reactive mAbs [15, 16, 17]. We therefore tested the reactivity of the seven mAbs against peptides equivalent to the first 15 (VP2N15), 30 (VP2N30) or 45 (VP2N45) amino acids of the N-terminus of VP2 from FMDV O1K (Fig.1a). The N-terminus of VP2 is known to be most highly conserved within the first 15 amino acids. The five mAbs (4D1, 1D6, 4A3, 5B2 and 5F10) identified as VP2-specific by Western blots also reacted strongly with the VP2 peptides in ELISA (Fig.1b). Among them, two mAbs (4A3 and 5B2) showed an equivalent reactivity with the three peptides, while the three remaining mAbs recognized the VP2N15 peptide with lower intensity (Fig.1b). The mAb 4A3 was taken forward for further characterisation. In particular, fine mapping using 15mer peptides with 10 amino acids overlaps (Fig.1a) showed that mAb 4A3 reacted with the 15mer peptide that corresponded to the N-terminus of VP2 and not with a 15mer starting at amino acid 6, confirming the presence of an epitope at the N-terminus of VP2 (Fig.1c). The mAb 4A3 specifically detected a protein band in western blot of the expected size for VP2 in cell lysates from infections with all 7 serotypes (Fig.1d) confirming that the epitope is linear, conserved and specific for VP2. MAb 4A3 also recognised virus infected cells when used as the primary antibody in indirect immunofluorescence microscopy of IBRS-2 cell cultures infected with type O FMDV (Fig.1e).

### VP2N peptides detect antibodies in sera from animals infected with all serotypes of FMDV

An indirect ELISA using peptides VP2N15, VP2N30 or VP2N45 was used to assess the presence of antibodies against the N-terminus of VP2 in a representative serum from an animal infected with type O FMDV. All three peptides captured antibodies, with the longer peptides producing a slightly higher signal (Fig.2a). A control peptide equivalent to a capsid sequence from the related picornavirus human rhinovirus gave a low signal consistent with background.

**Fig.2.**
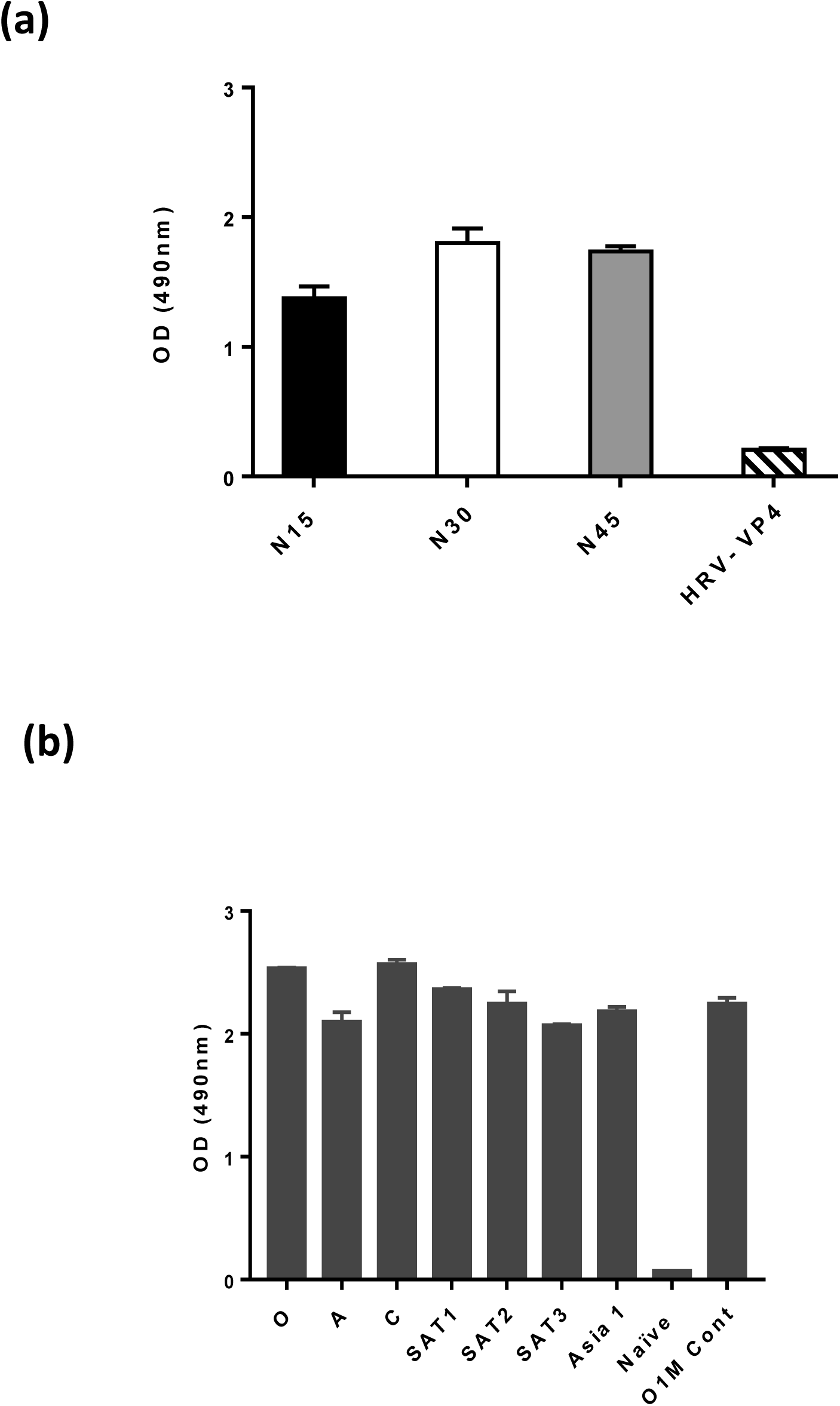
Sera from animals infected with any serotype of FMDV react with VP2 peptides. **(a)** Reactivity of serum from an animal experimentally infected with FMDV serotype O with peptides equivalent to the N-terminal 15 (N15), 30 (N30) or 45 (N45) amino acids of FMDV VP2, or the N-terminal 45 amino acids of human rhinovirus VP4 (HRV-VP4, negative control). **(b)** Reactivity of sera from animals vaccinated with vaccine strains of the seven serotypes with the FMDV VP2N-45 peptide.

The longer peptide VP2N45 was then used to test monovalent sera from different animals vaccinated against the seven serotypes of FMDV; this showed that the same peptide was able to detect antibodies against all the serotypes (Fig.2b).

### Development of a VP2 ELISA for universal detection of FMDV antibodies

A VP2 ELISA using peptide VP2N45 was developed using reference sera. The optimal concentration of peptide and dilution of sera to be used in the test was first evaluated by checkerboard titrations using bovine sera known to be negative or strongly positive or weakly positive for antibody by existing tests. The best signal to noise ratio (positive: negative) was obtained using a serum dilution of 1 in 100 and peptide concentration of 2μg/ml (Fig S.1). At these optimised conditions, the cut off for distinguishing between positive and negative signals was set as 0.4 OD units, calculated using the average value of three independent tests using the standard negative reference serum sample used by WRLFMD for routine FMDV diagnostics.

Using the optimized assay conditions, a collection of previously characterized serum samples was tested in triplicate and repeated twice independently, representing naïve cattle (n=100) and cattle vaccinated (n=38) or infected (n=34) with all seven serotypes of FMDV. The majority of vaccinated and infected (positive) samples gave a relatively strong signal (average absorbance value of 1.4) and the majority of naïve (negative) samples gave a relatively low signal below 0.4 (Fig.3a).

Seven negative sample exceeded the cut off of 0.4 OD units (ranging between 0.4 and 1.0 OD) and would be considered false positive, therefore producing a diagnostic specificity for the test of 93%. The signal for one positive sample (type A vaccinated) was below this cut off and would be considered a false negative in this test giving a sensitivity of 98.6%.

### Comparison of the VP2 ELISA with existing tests (VNT, LPBE, SPCE and PrioCHECK)

For the positive serum samples analyzed by VP2 ELISA in Fig.3a, pre-existing WRLFMD data generated using established diagnostic tests was accessed retrospectively and used to compare the performance of the VP2 ELISA. The pre-existing data was generated with four tests: VNT to quantitate neutralising antibodies, LPBE, SPCE and PrioCHECK to quantitate anti-capsid antibodies. The sensitivity of the VNT, LPBE and SPCE are dependent on close antigenic match between reagents used (virus/antigen and antibodies) and the serum sample being tested. Therefore, the data from VNT and LPBE were subdivided into groups carried out with homologous (same virus used to vaccinate or infect the animal) or heterologous (same serotype but strain different than those used to vaccinate or infect the animal) reagents. The data obtained with PrioCHECK kits was only available for samples from infections with serotypes O, A and Asia 1.

As mentioned above, the VP2 ELISA data (Fig.3a) contained a single false negative equivalent to a sensitivity of 98.6%. In comparison, the homologous VNT data (n=37) had no false negatives (sensitivity of 100%) while the heterologous VNT data (n=72) had a sensitivity of 73.2% (Fig.3b and Table 2). Similarly, the homologous LPBE data (n=30) had no false negatives (sensitivity of 100%) and the heterologous LPBE data (n=72) had several false negatives (sensitivity of 93.0%) (Fig.3c and Table 2). The SPCE data (n=72) had a single false negative (sensitivity of 98.6%) (Fig. 3d and Table 2) and the PrioCHECK data (n=29) had two false negatives (sensitivity of 93.1%) (Fig. 3d and Table 2).

**Table 2.**
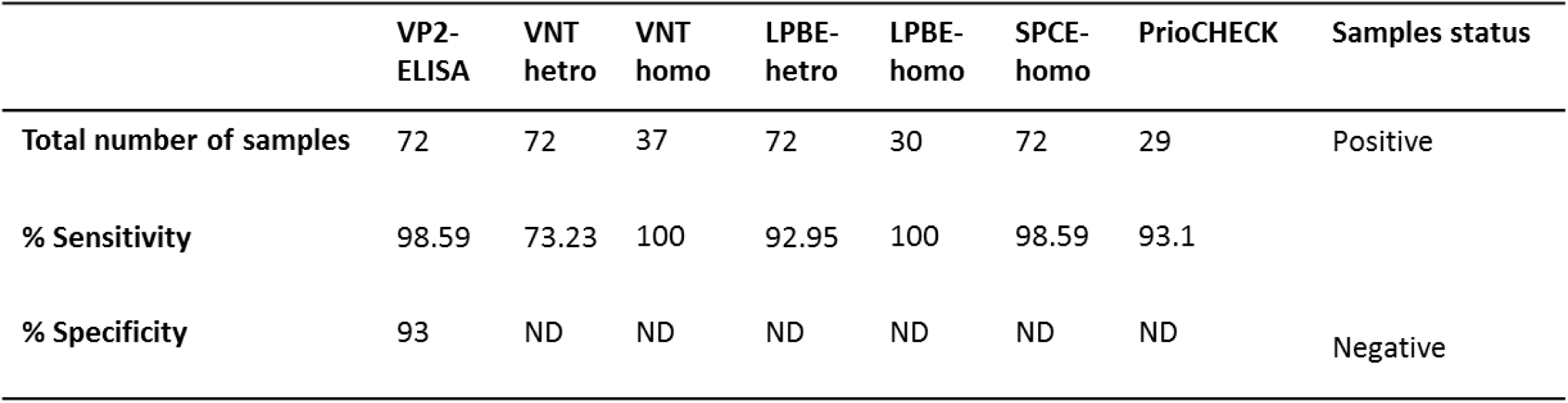
Comparative sensitivities of VP2 ELISA and other existing serological tests.

|  | VP2-ELISA | VNT<br>hetro | VNT<br>homo | LPBE-hetro | LPBE-homo | SPCE-homo | PrioCHECK | Samples status |
| --- | --- | --- | --- | --- | --- | --- | --- | --- |
| **Total number of samples** | 72 | 72 | 37 | 72 | 30 | 72 | 29 | Positive |
| **% Sensitivity** | 98.59 | 73.23 | 100 | 92.95 | 100 | 98.59 | 93.1 |  |
| **% Specificity** | 93 | ND | ND | ND | ND | ND | ND | Negative |

**Fig.3.**
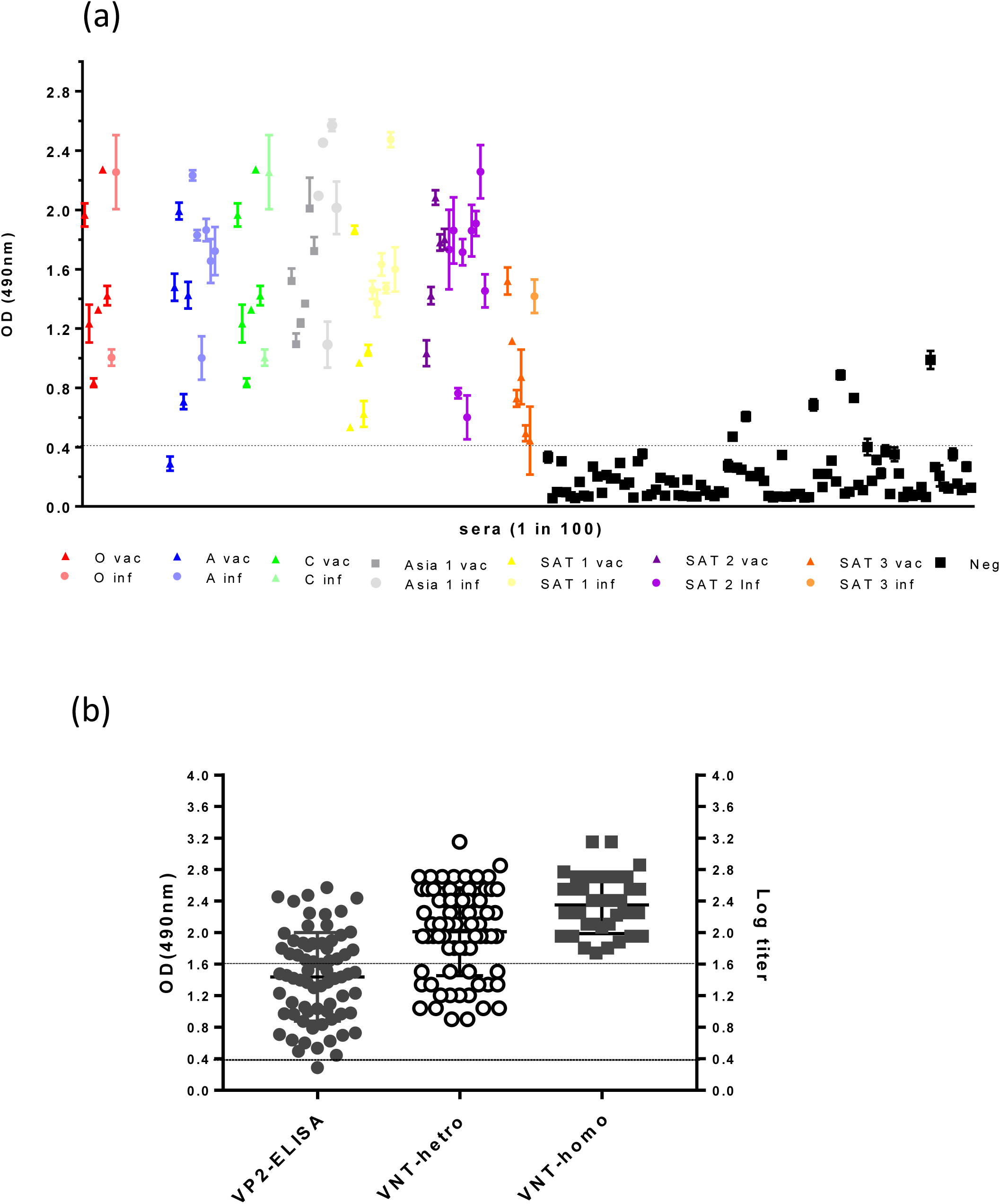
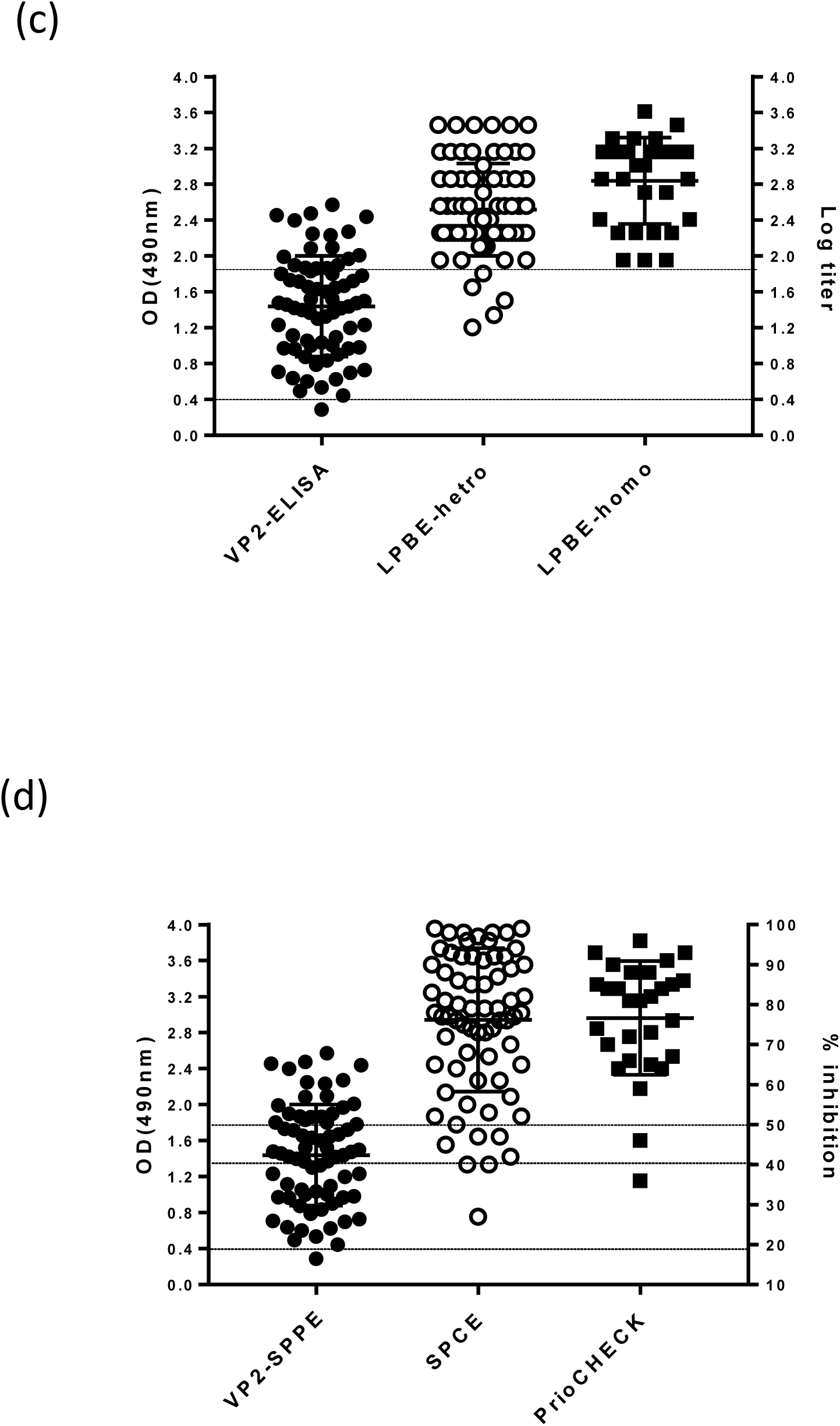
Testing reference negative and positive serum samples to detect the specificity and sensitivity of the assay. **(a)** Reactivity in VP2 ELISA (OD 490nm) of negative (black squares, n=100) and positive (circles are infected, triangles are vaccinated, serotypes represented by colours as indicated; n=72) reference sera. Peptide was at 2μg /ml and sera diluted 1 in 100. **(b)** Distribution plots comparing results of positive sera tested with VP2 ELISA (same as data in a; cut-off of 0.4 OD) and with homologous (n=37) and heterologous (n=72) VNT (cut-off = log titre 1.65). **(c)** Distribution plots comparing results of positive sera tested with VP2 ELISA (same as data in a; cut-off of 0.4 OD) with homologous (n=30) and heterologous (n=72) LPBE (cut-off =log titre 1.95). **(d)** Distribution plots comparing results of positive sera tested with VP2 ELISA (same as data in a; cut-off of 0.4 OD), with SPCE (n=30, cut-off =40% of inhibition) and PrioCHECK kits ELISA (n=29, cut-off =50% of inhibition)

The single false negative sample (A Eritrea 3/98-41dpv) in the VP2 ELISA was also a false negative in both the heterologous VNT (log titer =1.04) and heterologous LPBE (log titre=1.6), but was positive in homologous VNT (log titer of 2.06) and weakly positive in the SPCE (52 % inhibition) and PrioCHECK (65 % inhibition).

Overall these results show that the VP2 ELISA detected antibody to all serotypes and the OD values may provide an estimate of the level of antibodies. The sensitivity of the new test resulted equivalent to or better than PrioCHECK kits and SPCE; sensitivity was significantly higher than LPBE and VNT when such assays are carried out with heterologous reagents.

## Discussion

This study describes the development of a novel assay for the detection of antibodies against the FMDV capsid that can be used to test for seroconversion in infected or vaccinated animals. The benefits of this assay are that FMDV-specific SP antibodies from all seven serotypes can be detected without the requirement for individual specific antigen or antibody reagents that are required for existing tests such as VNT, LPBE, SPCE.

This assay targets a capsid epitope at the N-terminus of VP2 that exhibits high sequence conservation among all seven serotypes of FMDV. Cross-reactive mAbs and overlapping peptides were used to show that the minimum sequence required for this linear epitope was VP2-N 1-DKKTE-5. This is consistent with previous studies, where structures of the FMDV capsid suggested that the N-terminus of VP2 is an internal component but may be flexible allowing it to be present at the surface to contribute to antigenicity [22, 23, 24]. In addition, the production of monoclonal antibodies to VP2 N-terminus in response to immunisation with FMDV, suggested that capsid flexibility may expose some of the internal domains of the capsid proteins to the surface enabling them to become antigenic sites [15,16, 17]. It has also been reported that a purified recombinant 1AB (VP4/VP2) capsid protein was detected by antisera against all seven FMDV serotypes, indicating that the VP4/VP2 protein contained a highly conserved epitope. Peptides containing the VP2 N-terminal epitope were reactive with antibodies against all seven FMDV serotypes and one (VP2N45) was selected as the basis of a novel VP2 ELISA that was evaluated with a panel of reference sera from naïve (n=100), vaccinated (n=38) and infected (n=34) cattle, representative of all the seven FMDV serotypes. Results demonstrated that the VP2 ELISA detected antibody to all serotypes with a diagnostic specificity of 93% and sensitivity of 98.6%. The sensitivity of the new ELISA was equivalent to or better than existing tests, such as PrioCHECK kits and SPCE; sensitivity was significantly higher than LPBE and VNT carried out with heterologous reagents.

The VP2 ELISA is suitable for detection of antibodies against the capsid of FMDV either post vaccination or post infection. The capture antigen contains a universally conserved viral epitope that is expected to be present on any isolate of FMDV, this ensures that the VP2-ELISA is able to detect FMDV antibodies regardless of the viral strain. In contrast to the biological reagents necessary in many other ELISA, the VP2 capture antigen is a synthetic peptide, greatly facilitating standardisation, continuity of supply and reproducibility. More importantly, it does not require the optimisation and re-validation when serum from antigenic distant strains needs to be tested.

Serological testing is a suitable tool for FMD surveillance. Detection of NSP antibodies currently offers the advantages of a DIVA and cross-serotype test. However, the VP2 ELISA can be used as a complementary or confirmatory test to the NSP ELISA, which is especially useful in obtaining FMDV free status after an outbreak. As for the NSP ELISA, the VP2 ELISA can also be used as (1) a front-line sero-surveillance assay in areas which are normally free from FMD without vaccination, (2) for areas conducting surveillance to achieve free from vaccination status, and (3) at the point of import and export to confirm the freedom of animals from FMDV antibodies. The test may also provide a simple approach for evaluating vaccine efficacy in experimental and field trails, although additional studies would need to be carried out to determine the cut-off that correlates to protection.

In conclusion, the results suggest that the VP2 ELISA developed for the detection of antibodies to FMDV has potential applications as a rapid, simple and inexpensive test in the sero-diagnosis of FMDV and in sero-surveillance programmes. Further validation and standardisation will be required to confirm the potential benefits of the VP2 ELISA.

## Acknowledgments

This work was financially supported by BBSRC research grant BB/L004828/ and Defra project SE1129. The Pirbright Institute receives strategic support from the Biotechnology and Biological Sciences Research Council of the United Kingdom (projects BB/E/I/00007035 and BB/E/I/00007036)

## Supplementary figures and tables

**Fig.S1.**
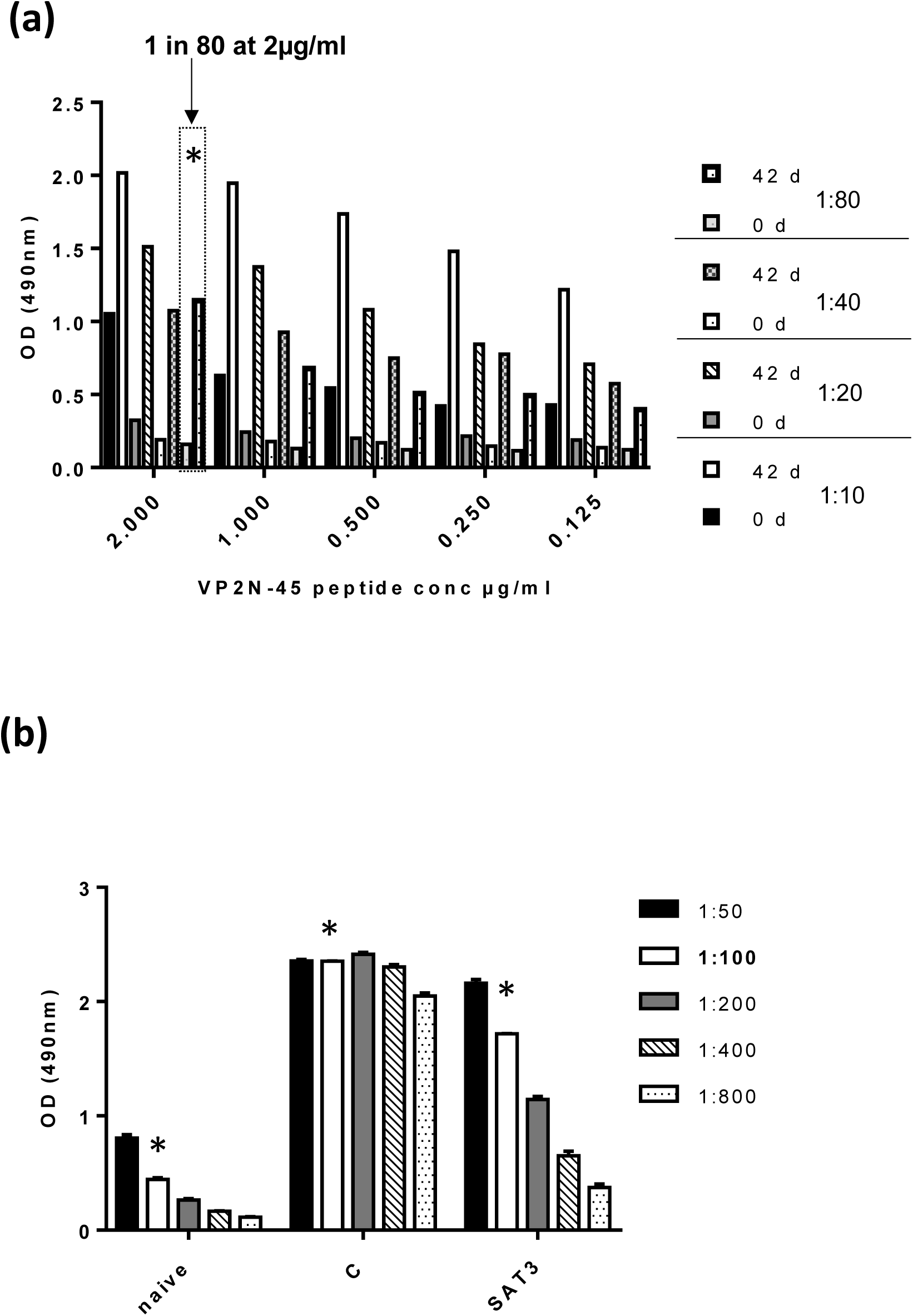
Optimisation of the peptide ELISA using different concentrations of peptide and dilution of the serum of serotype O from infected animal. **(a)** Checkerboard ELISA with negative (0 d) and positive (42 d) sera diluted from 1:10 to 1:80 (as shown in key) and with peptide concentration in the range 0.125-2μg/ml. The optimal conditions for signal to background are highlighted with a box (2μg/ml of peptide and 1 in 100 serum dilution). **(b)** Reactivity with VP2N45 peptide at 2μg/ml of different dilutions of a strong responder serum sample (type C) and a weak responder serum sample (type SAT3). The asterisk denotes the best conditions of peptide at 2μg/ml and sera diluted 1:100.

